# Pathway crosstalk enables cells to interpret TGF-β duration

**DOI:** 10.1101/134106

**Authors:** Jingyu Zhang, Xiao-Jun Tian, Yi-Jiun Chen, Weikang Wang, Simon Watkins, Jianhua Xing

## Abstract

The detection and transmission of the temporal quality of intracellular and extracellular signals is an essential cellular mechanism. It remains largely unexplored how cells interpret the duration information of a stimulus. In this paper, through an integrated quantitative and computational approach we demonstrate that crosstalk among multiple TGF-β activated pathways forms a relay from SMAD to GLI1 that initializes and maintains SNAILl expression, respectively. This transaction is smoothed and accelerated by another temporal switch from elevated cytosolic GSK3 enzymatic activity to reduced nuclear GSK3 enzymatic activity. The intertwined network places SNAIL1 as a key integrator of information from TGF-β signaling subsequently distributed through upstream divergent pathways; essentially cells generate a transient or sustained expression of SNAIL1 depending on TGF-β duration. Other signaling pathways may use similar network structure to encode duration information.

## Introduction

Cells live in a state of constant environmental flux and must reliably receive, decode, integrate and transmit information from extracellular signals such that response is appropriate^1-4^. Dysregulation of signal transduction networks leads to inappropriate transmission of signaling information, which may ultimately lead to pathologies such as cancer. Therefore a central problem in systems biology has been to untangle how quantitative information of cellular signals is encoded and decoded. In general cells respond to one or more properties of a stimulus, such as its identity, strength, rate of change, duration and even its temporal profile^5-11^. There are extensive studies on the dose-response curves to reveal how cells respond differentially to a signal with different strength. In comparison how cells respond to the temporal code of signals is less studies, and its physiological relevance receives much attention recently since most extracellular signals exist only transiently and cellular responses show dependence on signal duration^12-16^.

Transforming growth factor-β (TGF-β) is a secreted protein that regulates both transient and persistent cellular processes such as proliferation, morphogenesis, homeostasis, differentiation, and the epithelial-to-mesenchymal transition (EMT)^17-21^. Because it plays essential roles in developmental and normal physiological process, and its dysregulation is related to cancer, fibrosis, inflammation, Alzheimer’s disease and many other diseases, the TGF-β signaling pathway has been probed extensively as a potential pharmaceutical target^22,23^. Several quantitative studies have expanded our knowledge on how the TGF-β-SMAD signaling pathway transmits the duration and strength information of the signal^24-28^.

TGF-β can activate both Smad-dependent and multiple Smad-independent pathways, which then converge onto some downstream signaling elements. It is unclear how cells transmit and integrate information of the TGF-β signaling distributed among these parallel pathways. Addressing this question requires studies beyond the TGF-β-SMAD axis as in earlier work, where quantifying SMAD proteins serves as the fundamental readout^24-26^. Here we focused on expression of SNAIL1, which is such a downstream target and plays a key role in regulating a number of subsequent processes. Our results confirmed that crosstalk between the SMAD-dependent and independent pathways are key for cells to decode and transmit temporal and contextual information from TGF-β. We posit that the mechanism may be a central mechanistic signal transduction process as many signaling pathways share the network structure.

## Results

### Network analysis reveals a highly connected TGF-β signaling network

Through integrating the existing literature, we reconstructed an intertwined TGF-β-SNAIL1 network formed with SMAD-dependent and ‐independent pathways (Fig. S1). For further studies we then identified a coarse-grained network composed of a list of key molecular species (Fig. 1, and Supporting text for details). Along the canonical SMAD pathway, TGF-β leads to phosphorylation of SMAD2 and/or SMAD3 (pSMAD2/3), followed by nuclear entry after recruiting SMAD4 and forming the complex. The complex acts as a direct transcription factor for multiple downstream genes including SNAIL1 and I-SMAD^24,29^. I-SMAD functions as an inhibitor of pSMAD2/3, thus closes a negative feedback loop. TGF-β also activates GLI1, a key component of the Hedgehog pathway, both through transcriptional activation by pSMAD2/3, and through suppressing the enzymatic activity of glycogen synthase kinase 3 (GSK3). The latter is constitutively active on facilitating GLI1 and SNAIL1 protein degradation in untreated epithelial cells^30,31^, thus suppressing GSK3 is expected to lead to GLI1 and SNAIL1 protein accumulation. Therefore the network integrates multiple feed-forward loops that converge at the regulation of SNAIL 1 transcription. In the following sections we will examine the functional roles of individual pathways in the network using several human cell lines.

**Figure 1.**
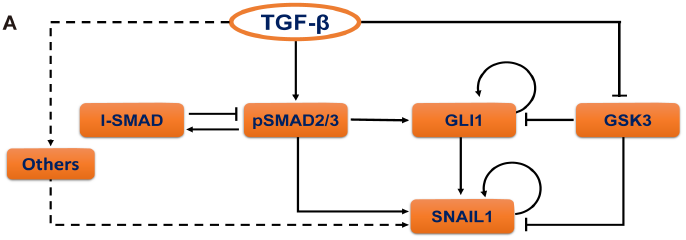
TGF-β induced signaling crosstalk network converges to SNAIL1. Reconstructed literature-based pathway crosstalk for TGF-β induced SNAIL1 expression. The node “Others” refer remaining SNAIL1 activation pathways that have minor contributions to the time window under study and thus are not explicitly treated.

### The canonical TGF-β/SMAD pathway initializes a transient wave of SNAIL1 expression

First we examined the TGF-β/SMAD/SNAIL1 pathway (Fig. 2A) by treating human MCF10A cells with recombinant human TGF-β 1, and performing multicolor immunofluorescence (IF) using antibodies directed against pSMAD2/3, SNAIL1. As expected from the pSMAD/I-SMAD negative feedback loop, pSMAD2/3 proteins accumulated in the nucleus transiently, peaking at 12 hours after TGF-β 1 treatment, followed by a decrease by 24 hours (Fig. 2B & C). We confirmed the transient pSMAD2/3 dynamics by sampling 1100-2600 cells at each time point (Fig. 2C). The result is also consistent with reports in the literature^24,26,32^. Nuclear SNAIL1 concentration rose concurrently with pSMAD2/3 (Fig. 2B & C), then there was a transient dip at 24 hours, followed by another increase then a persistant elevation for one week^33^.

**Figure 2.**
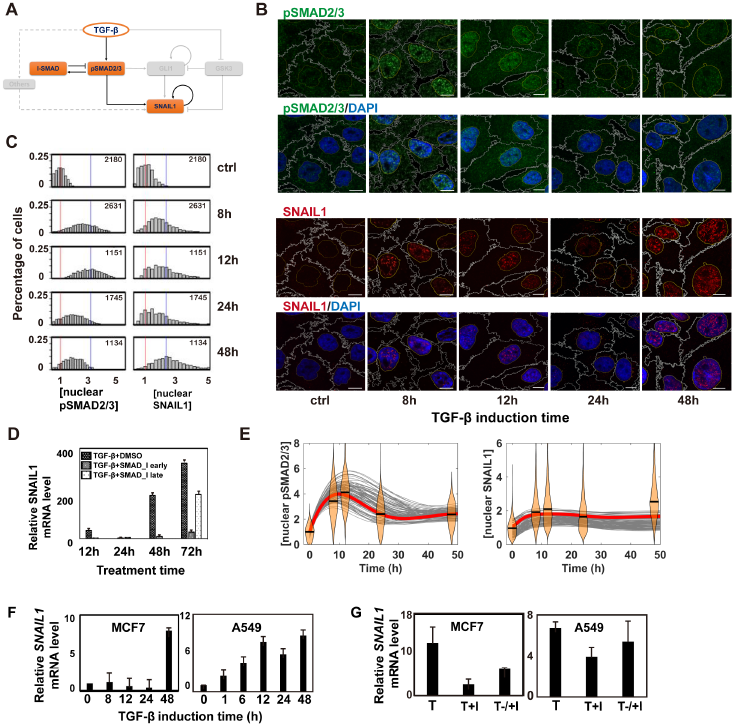
The SMAD proteins induce the first wave of SNAIL1. (A) Canonical SMAD-dependent pathway for TGF-β activation of SNAIL1 highlighted from the network in Fig 1. (B) Two-color immunofluorescence (IF) images of pSMAD2/3 and SNAIL1 of MCF10A cells induced by 4 ng/ml TGF-β1 at various time points. The scale bar is 10 *μm* and is the same for other IF images in this paper. (C) Distributions of nuclear pSMAD2/3 and SNAIL1 concentrations quantified from the IF images. Red vertical lines indicate the mean value of the distributions at time 0, and blue vertical lines represent that at 12 h (for pSMAD2/3) or at 48 h (for SNAIL1), respectively. The number in each figure panel is the number of randomly selected cells used for the analysis. Throughout the paper we report fold changes of concentration and amount relative to the mean basal value of the corresponding quantity. (D) Effects of early (added together with TGF-β) and late (48 h after adding TGF-β) pSMAD inhibition on the *SNAIL1* mRNA level in MCF10A cells. (E) Thorough parameter space search confirmed that with the model in panel A one can fit the pSMAD2/3 dynamics, but not the two-wave SNAIL1 dynamics. The experimental data are shown as violin plots with the medians given by black bars. Solid curves are computational results with parameter sets sampled from the Monte Carlo search, and the red curves are the best-fit results. (F) Fold change of *SNAIL1* mRNA levels in MCF7 and A549 cells measured with quantitative RT-PCR after TGF-β 1 treatment. (G) Fold change of *SNAIL1* mRNA levels measured with quantitative RT-PCR at 72 h after TGF-β1 (T) treatment. For early inhibition (T+I) the inhibitor was added at the time of starting TGF-β 1 treatment. For late inhibition (T-/+I) the inhibitor was added 48 h (for MCF7) and 24 h (for A549) after starting TGF-β 1 treatment, respectively. The inhibition results were compared to the TGF-β treatment (T) result at the same time point.

Next, we treated MCF10A cells with a SMAD2/3 active phosphorylation inhibitor LY2109761 in addition to TGF-β treatment (Fig. 2D). Without the inhibitor, the *SNAIL1* mRNA showed the two-wave dynamics consistent to that of the protein. When the inhibitor was added concurrently with TGF-β treatment, the *SNAIL1* mRNA level was reduced to ∽ 9% of that of the control experiment (without inhibitor) by day 3, indicating that indeed pSMAD2/3 are required for SNAIL 1 initial activation. However, the *SNAIL1* mRNA level remained ∽70% when the inhibitor was added 48 hours after initiation of TGF-β treatment (when nuclear pSMAD2/3 concentration has dropped to a minimum). Furthermore we constructed a mathematical model that contains only the TGF-β/SMAD/SNAIL1 pathway, and performed a thorough parameter space search using a multi-configuration Monte Carlo algorithm (Fig. S2B). The search revealed regions of the parameter space that quantitatively reproduced the transient pSMAD2/3 dynamics, but not the two-wave dynamics of SNAIL1 expression (Fig. 2E). This computational result further confirmed that pSMAD2/3 is less essential for the second wave of SNAIL1.

Furthermore, this SNAIL1 dynamics is not cell type dependent as equivalent two-wave dynamics were seen for *SNAIL1* mRNA in MCF7 and A549 cells (Fig 2F). Similar to that of MCF10A, it is more effective on inhibiting *SNAIL1* mRNA by adding LY2109761 together with TGF-β than later (Fig. 2G). In total, these results reveal that pSMAD2/3 is essential for the early phase of SNAIL1 activation, but is less important for the secondary phase elevation and persistence of SNAIL1 expression/localization.

### GLI1 contributes to activating the second wave of SNAIL1

The regulatory network suggests that GLI1 may be responsible for the second wave of SNAIL1 (Fig. 3A). To test this hypothesis, we performed microscopy studies of SNAIL1-GLI1 using MCF10A cells. The distribution of SNAIL1 found in this study (Fig. S3A) was consistent with those from the pSMAD2/3-SNAIL 1 studies. Elevated and sustained expression of GLI1 under TGF-β treatment (Fig. 3B & C) was clearly evident. More interestingly GLI1 also showed an unexpected multi-phasic dynamic. Around 8 hours after TGF-β treatment, cytosolic GLI1 concentration started to increase. At 12 hours when SMAD activities decreased toward basal levels there was a clear accumulation of GLI1 in the nucleus, which continued to increase through day 2. Notably, at this time point cells expressing a high level of nuclear SNAIL1 consistentlyshowed high intranuclear concentrationsGLI1(Fig.S3A). Expanding the mathematical model of the network to Fig. 2A also reproduced the temporal dynamics of pSAMD2/3 and SNAIL1 (Figure S3B), supporting the role of GLI1 as the activator of the second waveofSNAIL1.

**Figure 3.**
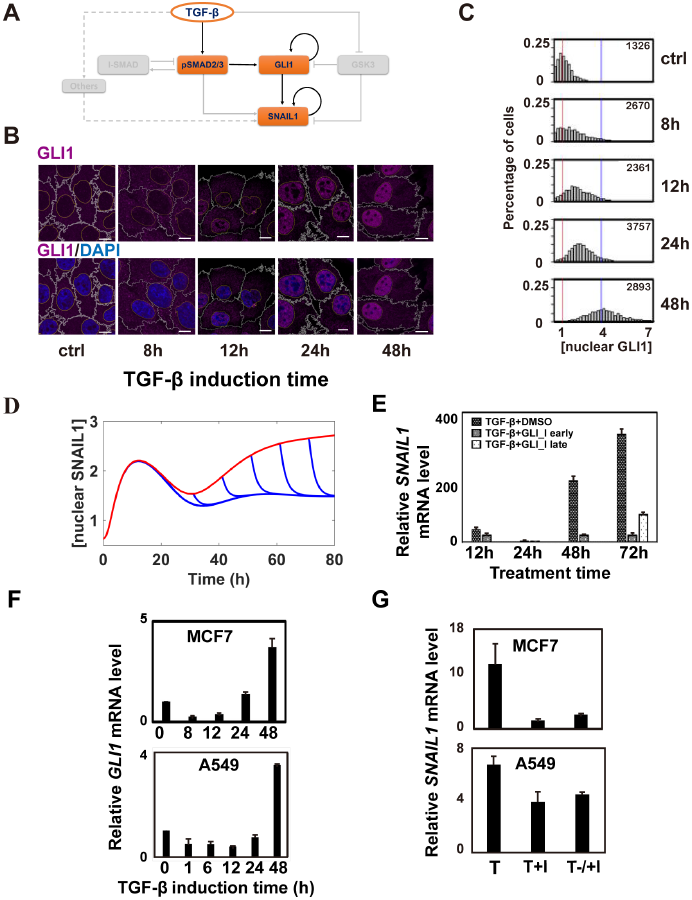
GLI1 is a major contributor to activate the second wave of SNAIL1 expression. (A) TGF-β activates the GLI1/SNAIL1 module partly through pSMAD2/3. (B) IF images on protein levels of GLI1 (in the free form). Red and blue vertical lines indicate the mean values of the distributions at time 0 and at 48 h, respectively. (C) Distributions of nuclear GLI1 concentrations quantified from the IF images. (D) Predicted outcome of adding GLI1 inhibitor at different time after TGF-β 1 treatment (blue lines). The red line is the predicted dynamics without the inhibitor. (E) Experimental validation of the results for early (added together with TGF-β) and late (48 h after adding TGF-β) GLI1 inhibition on the *SNAIL1* mRNA level in MCF10A cells. (F) Fold change of *GLI1* mRNA levels measured with quantitative RT-PCR at different time points after combined TGF-β 1 treatment in MCF7 or A549 cells. (G) Fold change of *SNAIL1* mRNA levels measured with quantitative RT-PCR at 72 h after combined TGF-β 1 and GLI1 inhibitor GANT61 treatment in MCF7 or A549 cells. For early inhibition (T+I) the inhibitor was added at the time of starting TGF-β 1 treatment. For late inhibition (T-/+I) the inhibitor was added 48 h (for MCF7) and 24 h (for A549) after starting TGF-β 1 treatment, respectively. TGF-β treatment group (T) is shown as a positive control.

If GLI1 is involved in the maintenance of SNAIL1 expression subsequent to the drop in pSMAD2/3 concentration, it is reasonable to predict (Fig. 3D) that inhibiting GLI1 activity, either at the onset of or at some subsequent time after TGF-β treatment, would have minimal effect on the initial wave of SNAIL1 expression since the latter is caused by pSMAD2/3. However, it would eliminate the second wave of SNAIL1 expression. Indeed this was observed when the GLI1 inhibitor GANT61 was added together with TGF-β at the beginning of the experiment resulting in a reduction of the *SNAIL1* mRNA level to be 55% (at 12 h and 24 h), 12% (at 48 h) and 7% (at 72 h) compared to those without inhibition at the corresponding time points (Fig. 3E). In another experiment adding the inhibitor 48 hours after TGF-β treatment also reduced the mRNA level measured at 72 h to be 25% (Fig. 3E). These results are qualitatively different from those with the SMAD inhibitor (Fig. 2D).

To confirm that GLI1 activation is not restricted to the MCF10A cell line, we also examined MCF7 and A549 cells with TGF-β treatment, and observed similar increased and sustained GLI1 expression (Fig. 3F). Furthermore, early and late GLI1 inhibition lead to a reduction of the *SNAIL1* mRNA level to be 13% and 22% for MCF7 cells, and to a less extent of 57% and 66% for A549 cells, respectively (Fig. 3G). Additionally, increased GLI1 expression after TGF-β treatment has been found for multiple liver cancer cell lines^34^. *In toto* these results support the role of GLI1 as a signaling relay from pSMAD2/3 to SNAIL1.

### GSK3 in a phosphorylation form with augmented enzymatic activity accumulates at endoplasmic reticulum and Golgi apparatus

Next, we hypothesized that GSK3 is fundamental to the observed multi-phasic GLI1 dynamic (Fig. 3B). Most published studies suggest that GSK3 is constitutively active in untreated cells, facilitating degradation of SNAIL1 and GLI1; TGF-β treatment leads to GSK3 phosphorylation and inactivation, which leads to an accumulation of SNAIL1 and GLI1^35,36^.

Initially we tested whether the above mechanism is sufficient to explain the multi-phasic GLI1 dynamics. We treated MCF10A cells in the absence of TGF-β with a GSK3 activity inhibitor. Given the above mechanism, one should expect the GSK3 inhibitor to promote both GLI1 and SNAIL1. In our experiment, SNAIL1 did increase, but there was no change in GLI1 expression in either nucleus or cytoplasm (Fig. 4A), suggesting additional signaling mechanisms may be involved.

**Figure 4.**
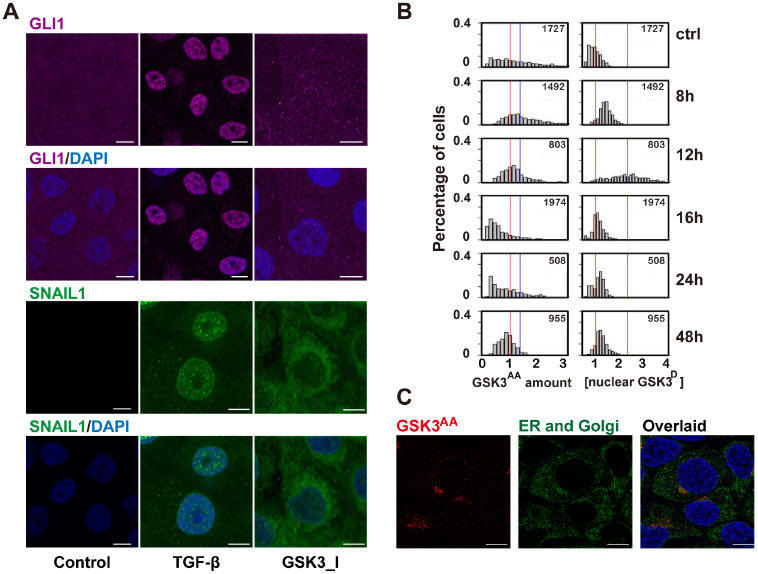
TGF-β induced temporal switch between active and inhibitive phosphorylation forms of GSK3 proteins. (A) IF images showed that inhibiting GSK3 enzymatic activity alone increased SNAIL1 accumulation but did not recapitulate TGF-β induced GLI1 nuclear translocation. (B) Quantification of the IF images of MCF10A cells at different time points after TGF-β treatment. Red vertical lines indicate the mean value of the distributions at time 0, and blue vertical lines represent that at 8 h (for GSK3^AA^) or at 12 h (for GSK3^D^), respectively. (C) IF images showing GSK3^AA^ localization at the endoplasmic reticulum center (ERC).

Besides the inhibitory serine phosphorylation, the literature suggests that tyrosine (Y279 in GSK-3α and Y216 in GSK-3β) phosphorylation leads to augmented enzymatic activity of GSK3^37^. As a convenience when discussing the three forms of GSK3, we refer the enzymatically active unphosphorylated form and the more active tyrosine phosphorylated form as “GSK3^A^” and “GSK3^AA^”, respectively, and the inactive serine phosphorylation form as “GSK3^D^”. Also we reserve “GSK3” for the total GSK3. As expected, microscopy studies showed an increased concentration of GSK3^D^ peaking around 12 hours after TGF-β treatment (Fig. 4B, Fig. S4A). Large cell-to-cell variations in the concentration of GSK3^D^ were observed, however, the abundance of cytosolic and nuclear GSK3^D^ were essentially equivalent (the expression ratio was close to one) for cells without TGF-β treatment (Fig. S4B). This observation corroborates earlier report that the serine phosphorylation does not affect GSK3 nuclear location^38^. TGF-β treatment led to transient deviation of this ratio from equivalence, reflecting additional active and dynamic regulation of GSK3 including covalent modification, location and protein stability. Specifically prior to inhibitory serine phosphorylation we observed transient GSK3^AA^ accumulation in the perinuclear region peaking at eight hours (Fig. 4B, Fig. S4A). Close examination of higher magnification confocal images revealed that the GSK3^AA^ formed clusters in the endoplasmic reticulum (ER) and Golgi apparatus, but not associated with actin filaments (Fig. 4C, Movie 1 & 2). Given that a function of active GSK3 is to modify target proteins post-translationally, our observation suggests an unreported role for GSK3^AA^ accumulating at the ER and Golgi apparatus is to modify newly synthesized proteins before their release to the cytosol. Specifically previous studies showed that in mammalian cells a scaffold protein SUFU binds to GLI to form an inhibitory complex, and SUFU phosphorylation by GSK3β prevents the complex formation, exposes the GLI1 nuclear localization sequence^39^, which explains the observed increase of free GLI1 in the cytosol followed by nuclear translocation (Fig. 3B). Since the two phosphorylation forms, GSK3^AA^ and GSK3^D^, coexist within single cells at defined time points, we performed co-immunoprecipitation and found that the probability of having the two GSK3 phosphorylation forms in one molecule was undetectable (Fig. S4C).

Contrary to our observation that TGF-β regulates GSK3^AA^ dynamics, other studies posit that GSK3^AA^ is not regulated by external cues^40^. To resolve this paradox, we measured the relative amount of different GSK3 forms through silver staining (Fig. S4D). Among the three forms, the overall percentage of GSK3^D^ increased from a basal level of 37% to 65% at 12 h after TGF-β treatment. In contrast, only a small fraction of GSK3 molecules assumed the GSK3^AA^ form and its overall abundance was stable over time (from ∽10% basal level to ∽13% at 8 h then back to ∽10% at 12 h after TGF-β treatment). Essentially levels of GSK3^AA^ did not change in abundance but did change in localizations (homing to the ER and Golgi apparatus) to form a high local concentration, which imbue an important role in TGF-β signaling.

### A temporal and compartment switch from active to inhibitory GSK3 phosphorylation smoothens the SMAD-GLI1 relay and reduces cell-to-cell heterogeneity on GLI1 activation

Based on the above results, we constructed an expanded network for TGF-β induced SNAIL1 expression, which integrates a role for GSK3 and its switching among the three phosphorylation forms as well as a functional role due to its intracellular redistribution (Fig. 5A). The model reproduces the multiphasic dynamics of GLI1 as well as that of pSMAD2/3 and SNAIL1 (Fig. S5A).

**Figure 5.**
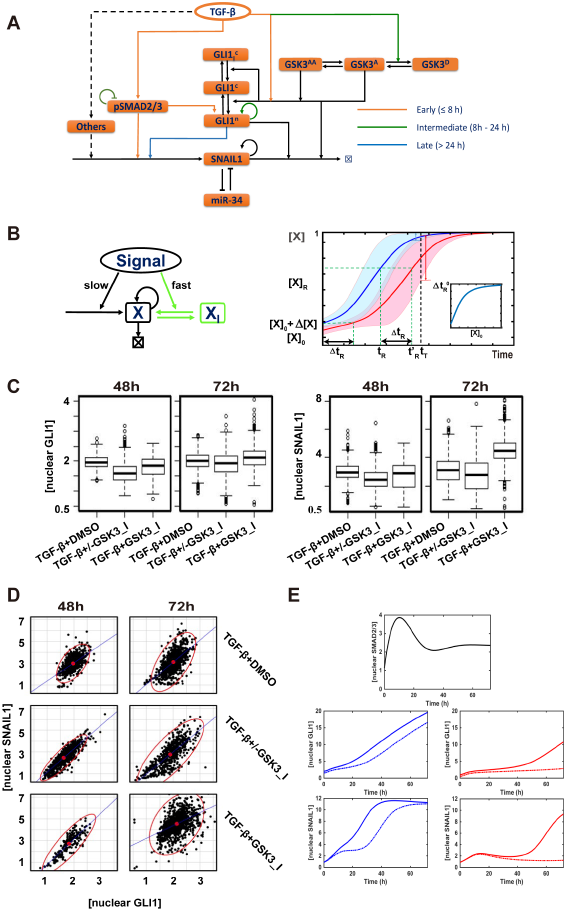
The GSK3 phosphorylation switch smoothens the SMAD-GLI1 relay. (A) Proposed expanded network for TGF-β induced SNAIL1 expression. (B) *left:* Schematic of a generic positive feedback loop network. Also shown in green is an additional reservoir of the molecules in inactive form (XI) that can convert quickly into the active form (X) upon stimulation. *Right:* The response time tRis sensitive to the initial concentration, [X]_0_v.s. [X]_0_ + Δ[X]_0_. The inlet figure shows the dependence of ΔtR on [X]_0_ with Δ[X]_0_ fixed. (C) Box plots of GSK3 inhibition experimental data. (D) Scattered plots of GSK3 inhibition experimental data. Red points are the center of the scattered plots and each ellipse encloses 97.5% of the data points. Both were drawn with the R package, *car::data.ellipse.* (E) Computational simulation of SMAD, SNAIL1 and GLI1 behavior with (solid line) or without (dotted line) initial boosting in cells with high basal GLI1 level (left panels) or low basal GLI1 (right panels).

To understand the function of the early nuclear accumulation of GLI1 induced by GSK3^AA^ , it is important to recognize that GLI1 has a positive-feedback loop, and this network motif (Fig. 5B, left panel) has characteristic sigmoidal shaped temporal dynamics, with the substrate concentration increasing slowly at first then accelerating with time until it approaches saturation (Fig. 5B, right panel, red curve). The response time, tR, defined as the time taken to reach a target concentration value [X]_R_, is highly sensitive to initial substrate concentration [X]_0_: in fact a slight increase in the initial concentration, Δ[X], will significantly shorten the response time (Fig. 5B, right panel, blue curve), and for a fixed Δ[X] a greater acceleration is seen in cells with lower initial concentrations (Fig. 5B, right panel, inlet figure). Consequently despite variations of their initial concentration of X, most cells within a population can reach [X]_R_ by a targeted time point t_T_ in a series of temporally regulated events such as cell differentiation and immune response. The needed concentration boost may be effected by conversion of preformed molecules from an inactive form into an active form (Fig. 5B, left panel, the part of the network in green). Indeed, many examples of this modified feedback loop motif exist. Figure S5B gives some examples involving members of intrinsically disordered proteins and inhibitors of DNA binding proteins, β-catenin and the STING motif for immune responses. In the present scenario the accelerated GLI1 dynamic ensures sufficient accumulation of GLI1 before nuclear pSMAD2/3 level decreases, essentially analogous to a relay race when the first runner can only release the baton after the second runner has grabbed it. Later when the GLI1 and SNAIL1 concentrations start to increase, the GSK3^A^→GSK3^D^ conversion became necessary to reduce the rates of their degradation catalyzed by active GSK3. Interestingly, this conversion takes place concurrently with maximal concentration of nuclear pSMAD2/3, which activates GLI1 and SNAIL1 transcription. Furthermore, the small initial concentration boost does not affect another major function of the positive feedback loop, which is to robustly buffer fluctuations in signal strengths via bistable dynamics (Fig. S5C).

To test the functional roles of GSK3 suggested above, we performed a series of GSK3 activity inhibition experiments. First, we pretreated MCF10A cells with GSK3 inhibitor SB216763, washed out the inhibitor then added TGF-β1 (Fig. S5D). We predicted that the treatment would slow down GLI1 nuclear accumulation, and at later times decrease the overall increase of GLI1 and SNAIL1 compared to cells without GSK3 inhibitor. Indeed this was observed (Fig. 5C, TGF-β +/− GSK3_I). More interestingly, the scatter plots (Fig. 5D) show the distributions with and without the inhibitor are similar in cells with high GLI1, but in the presence of inhibitor there is a population of non-responsive cells with low GLI1 and SNAIL1. This observation is consistent with model predictions that the GSK3-induced boost of initial GLI1 concentration leads an acceleration in the GLI1 and SNAIL1 dynamics, and this boost is more important for cells with lower level of initial nuclear GLI1 (Fig. 5E). In a separate experiment (Fig. S5E), we did not wash out GSK3 inhibitor while adding TGF-β. In this case the inhibitor had opposite effects on the GLI1/SNAIL1 protein concentration: it slowed down the initial release and translocation of GLI1 needed to accelerate the GLI1 accumulation, but also decreased GLI1 and SNAIL 1 degradation that becomes pre-eminent when the proteins were present at high levels. Compared to the samples grown in the absence of the GSK3 inhibitor, we also observed slower and more scattered GLI1 nuclear accumulation and SNAIL1 increase on day 2, but by day 3 the overall levels of GLI1 and SNAIL1 were actually higher than the case without the inhibitor (Fig. 5D, TGF-β + GSK3_I).

### The SMAD-GLI1 relay increases the network information capacity and leads to differential response to TGF-β duration

Our results show that TGF-β 1 signaling is effected through pSMAD2/3 directly with fast pulsed dynamics concurrently with a relay through GLI1 which has a much slower dynamics. The signaling ported by these two channels converges on SNAIL1 with a resultant two-wave expression pattern. To further dissect the potential functional interactions between these two pathways, we performed mathematical modeling and predicted that the two distinct dynamics allows cells to respond to TGF-β differentially depending on stimulus duration (Fig. 6A). Short pulses of TGF-β only activate pSMAD2/3 and the first wave of transient SNAIL1 expression. When the signal duration is longer than a defined threshold value, activation of GLI1 will lead to the observed second wave of SNAIL 1 expression. We confirmed the predictions with MCF10A cells (Fig. 6B). Both TGF-β1 pulses with duration of two hours and eight hours activated pSMAD2/3 and the first wave of SNAIL1 expressions. However, only the eight-hour but not the two-hour pulse activated sustained GLI1 and the second wave of SNAIL1 expression, similar to those with continuous TGF-β 1 treatment.

**Figure 6.**
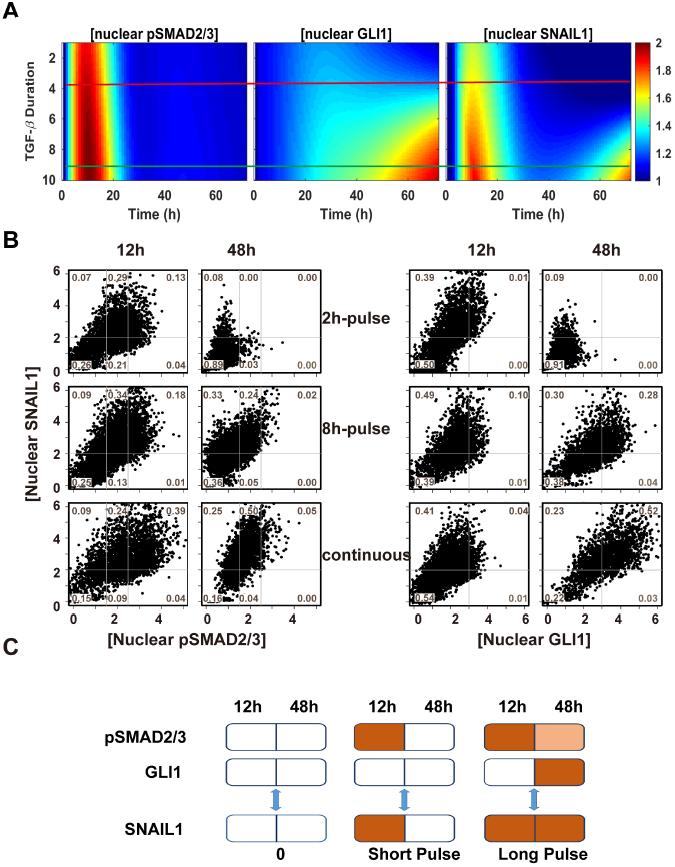
The TGF-β-SNAIL1 network permits detection of TGF-β duration and differential responses. (A) Model predictions that the network generates one or two waves of SNAIL1 depending on TGF-β duration. (B) Single cell protein concentrations quantified from IF images of cells under pulsed and continuous TGF-β treatments. The solid lines divide the space into coarse-grained states with respect to the corresponding mean values without TGF-β treatments (= 1). (C) Schematics of how cells encode information of TGF-β duration through a temporally ordered state space.

Clearly cellular responses have different temporal profiles depending on the TGF-β duration, and one can use the information theory to quantify their information content^9,10^. In this study we utilized a more intuitive understanding of network function from an information encoding viewpoint. Consider the pSMAD complex, which has three coarse-grained states, High (H), Medium (M), and Low (L), and each of GLI1 and SNAIL1 has two states, H and L (Fig. 6C). Then one can use three 4-element states, (L, L; L, L), (H, L; L, L), (H, M; L, H) to roughly describe the case without TGF-β and the 8 hour and 12 hour pulse results in Fig. 6B, where each number in a state represents in the order the 12 h and 48 h concentrations of pSMAD2/3 and GLI1, respectively. The three states are part of a temporally ordered state space, and encode information of TGF-β duration roughly as not detectable, short, and long. The same information is encoded by the SNAIL1 dynamics as (L, L), (H, L), and (H, H), reflecting SNAIL1 as an information integrator of the two converging pathways.

Further modeling suggests that components in the network function cooperatively to encode the TGF-β information (Fig. S6A). Increasing or decreasing the nuclear GSK3 enzymatic activity tunes the system to generate the second SNAIL 1 wave with a higher or lower threshold of TGF-β duration, respectively, while changing the cytosol GSK3 enzymatic activity has the opposite effect. Upregulation of GLI1, or downregulation of I-SMAD, both of which have been observed in various cancer cells, also decrease the threshold for generating the second SNAIL1 wave. Therefore cells of different types can share the same network structure, but fine-tune their context-dependent responses by varying some dynamic parameters, and for a specific type of cells dysregulation of any of the signaling network components may lead to misinterpretation of the quantitative information of TGF-β signal.

We have shown that the SNAIL 1 dynamics is TGF-β 1 duration dependent. To further confirm that cells respond differentially to TGF-β1 with different duration, we measured the mRNA levels of another four genes, all of which respond to TGF-β1 (Fig. S6B)^41,42^. Gene *FN1* codes for the cell motility related protein fibronectin. Its expression is activated even by the 2h TGF-β1 pulse, and increases with longer TGF-β1. Gene *CTGF,* whose product is an extracellular matrix protein and related to cell motility, is activated at similar extent by both 2h and 8h TGF-β1 pulses, and its expression level increases by additional 14 folds with continuous TGF-β1 treatment. Expressions of genes *MMP2 and CLDN4,* coding proteins related to mesenchymal extracellular hallmark and cell migration, increase only slightly (less than two folds) with either 2h or 8h TGF-β1 pulse, compared to the more significant change under continuous TGF-β1 treatment. Therefore, these downstream genes also show differential expression patterns depending on TGF-β1 duration, and cells activate different response programs correspondingly.

## Discussion

TGF-β is a multifunctional cytokine that can induce a plethora of different and mutually exclusive cellular responses. A significant open question is how cells interpret various features of the signal and make the cell fate decision. TGF-β can activate a number of pathways interconnected with multiple crosstalk points. Our studies reveal that this interconnection is essential such that components of the network can function coordinately and appropriately to interpret the temporal (time and duration) information from TGF-β.

### pSMADs are major inducers for the first wave of SNAIL1 expression

The two-wave dynamic of TGF-β-induced SNAIL1 expression has been observed in several cellular systems^43,44^, supporting the underlying relay mechanism discovered in this work. The first wave is fundamentally induced by pSMAD2/3, as evidenced from our SMAD inhibition experiments, and similarity between the dynamics of pSMAD2/3 and the first wave of SNAIL1. SNAIL1 may act as cofactor of pSMADs to induce other early response genes^45^. At later times the nuclear concentrations of pSMAD2/3 decrease though continue to contribute to SNAIL 1 activation at a lower level.

### GLI1 is a signaling hub for multiple pathways and temporal checkpoint for activating second-wave of sustained SNAIL1 expression

GLI protein has been traditionally attributed to the canonical Hedgehog pathway. Here we show that TGF-β induction of GLI1 relays the signal to induce SNAIL1. Consistent with the present study, Dennler et al. reported SMAD3-dependent induction of the GLI family by TGF-β both in multiple cultured cell lines, and in transgenic mice overexpressing TGF-β^30^ Many other signals such as PGF, EGF can also activate the GLI family, and a GLI code has been proposed to integrate input from different pathways and lead to context-dependent differential responses^46^. Our results confirmed this role of GLI1 as an intermediate information integrator and transmitter, and suggest that TGF-β must act above a threshold value of duration to activate the second wave of SNAIL1. This temporal checkpoint prevents spurious SNAIL1 activation and subsequent major cellular fate changes.

### GSK3 fine-tunes the threshold of the GLI1 checkpoint and synchronizes responses of a population of cells

The functional switch from pSMAD2/3 to GLI1 relays information from TGF-β signaling beyond the initial induction of SNAIL 1, and this relay is facilitated by a second relay from the active to the inactive phosphorylation form of GSK3 proteins. Active regulation of the abundance and nuclear location of GSK3^AA^ form has been observed in neurons^47^. In contrast to these earlier reports we observed an accumulation of GSK3^AA^ in the ER and Golgi apparatus. Mechanistically this may be caused by redistribution of cytosolic GSK3^AA^, or a simple accumulation of *de novo* synthesized and phosphorylated GSK3 proteins. The overall consequence is an increase in local GSK3 enzymatic activity, which forms part of the GSK3 switch that smooths the pSMAD2/3-GLI1 transition and the duration threshold of TGF-β pulse that generates the second wave of SNAIL1.

This seemingly simple process, which accelerates the response time through transient and minor increases in the initial concentration of a molecular species subject to positive feedback control, may have profound biological functions. Positive feedback loops are ubiquitous in cellular regulation, with a major function to filter both the strength and temporal fluctuations of stimulating signals and to prevent inadvertent cell fate change. This network, however, may have an inherently slow response time, and the response is highly sensitive to initial conditions that lead to large cell-to-cell variation of temporal dynamics. This variation and slow dynamic may be problematic for processes such as neural crest formation and wound healing where precise and synchronized temporal control is crucial for generating collective responses of multiple cells. Transient increases in initial protein concentrations of the GSK3 module may solve the seemingly incompatible requirements for the motif on robustness against fluctuations as well as fast and synchronized responses. It assures that despite a possible broad distribution of basal expression levels of the protein, cells are activated within a designated period of time at the presence of persistent activation signal, without scarifying the filtering function of the feedback loop.

### Cells use a temporally ordered state space formed by a composite network to increase information transfer capacity

Cells constantly encounter TGF-β signals with different strengths and duration, and must respond accordingly. It is well documented that biological networks reliably transmit information about the extracellular environment despite intrinsic and extrinsic noise in a subtle and functional way. However, quantitative analyses using information theory reveal that the dynamic of each individual readout is quite coarse with one or few bits^9,10^. This is a paradox. However, our results suggest that cells use multiple readouts to generate a temporally ordered state space with an expanded capacity to encode signal information and generate a far more subtle response system. For example, the SMAD motif has a refractory period due to the negative feedback loop and thus can accurately encode the duration information of TGF-β only within a limited temporal range, then the GLI1 motif encodes information of longer TGF-β duration, which then saturates. This temporally ordered state network may be further expanded, such that the SNAIL1 motif itself possibly encodes information of longer TGF-β duration and relays to other transcription factors such as TWIST and ZEB, and lead to stepwise transition from the epithelial to the mesenchymal phenotype depending on the TGF-β duration^33^.

As with other signaling process, TGF-β signaling is context dependent, and the dynamic and regulatory network vary between cell types^25,48^. For the three cell lines we examined our results identify GLI1 as a major relaying factor for the TGF-β signaling. The inhibition experiments show that other possible peripheral links have minor contributions to SNAIL1 activation, while their weights may grow at time later than we examined. Consequently the present work has focused on the early event of TGF-β activation of SNAIL1, which is with 72 h for MCF10A cells. Nevertheless, the relay mechanism and the corresponding network structure identified here can be general for transmitting quantitative information of TGF-β and other signals. It is typical that an extracellular signal is transmitted through a canonical pathway with negative feedbacks and multiple non-canonical pathways, and these pathways crosstalk at multiple points, and Fig. S6C gives some examples including IL-12, DNA double strand breaking, and LPS. Therefore the mechanism revealed in this work is likely beyond TGF-β signaling.

### Network temporal dynamics is a key for effective pharmaceutical intervention

Upregulation of GLI1, and GSK3 and the responsive SMAD family has been reported in pathological tissues of fibrosis^49^ and cancer^46^, and all three have been considered as potential drug targets. The present study emphasizes that in cell signaling timing is fundamental to function. The same network structure may generate drastically different dynamics with different parameters, as observed for different cell types. Consequently, effective biomedical intervention needs to take into account the network dynamics. We have already demonstrated that adding the inhibitors at different stages of TGF-β induction can be either effective versus not effective on reducing SNAIL1 (by inhibiting pSMAD2/3), both effective (by inhibiting GLI1), and even opposite (by inhibiting GSK3). Actually, one may even exploit this dynamic specificity for precisely targeting certain group of cells while reducing undesired side effects on other cell types.

In summary through integrated quantitative measurements and mathematical modeling we provided a mechanistic explanation for the long-time puzzle of how cells read TGF-β signals. Several uncovered specific mechanisms, such as expanding information transmission capacity through signal relaying, and reducing response times of positive feedback loops by increasing initial protein concentrations, may be general design principles for signal transduction.

## Materials and methods

### Cell Culture

MCF10A cells were purchased from the American Type Culture Collection (ATCC) and were cultured in DMEM/F12 (1:1) medium (Gibco) with 5% horse serum (Gibco), 100 μg/ml of human epidermal growth factor (PeproTech), 10 mg/ml of insulin (Sigma), 10 mg/ml of hydrocortisone (Sigma), 0.5 mg/ml of cholera toxin (Sigma), and 1x penicillin-streptomycin (Gibco). MCF7 cells were purchased from ATCC and cultured in EMEM medium (Gibco) with 10% FBS (Gibco), 10 mg/ml of insulin, and 1x penicillin-streptomycin. A549 cells were purchased from ATCC and were cultured in F12 medium (Corning) with 10% FBS and 1x penicillin-streptomycin. All cells were incubated at 37 °C with 5% CO2.

### TGF-β induce and inhibitor treatment

Cells for TGF-β induction and inhibitor treatment were seeded at ∽60-70% confluence without serum starvation. For TGF-β treatment, 4 ng/ml human recombinant TGF-β1 (Cell signaling) was added into culture medium directly. For inhibition experiment, 4 μM of LY2109761 (Selleckchem), 20 *μM* of GANT61 (Selleckchem), and 10 μM of SB216763 (Selleckchem) were used to inhibit SMAD, GLI and GSK3, respectively. The medium was changed every day during treatment to keep the reagent concentration constantly. For reproducibility, we used cells within 10^th^ -15^th^ generations, same patches of reagents, serum, and tried to perform each group of experiments (e.g., those in Fig. 2C) together.

### Immunofluorescence Microscopy and Data Analysis

Cells were seeded on four-well glass-bottom petri dishes at ∽60% confluence overnight and treated with reagents (TGF-β 1 and/or inhibitors). Three independent experiments were performed in every treatment. At designated time points, cells were harvested and stained with specific antibodies following procedure modified from the protocols at the Center of Biological Imaging (CBI) in the University of Pittsburgh. In general, cells were washed with DPBS for five minutes for three times followed by 4% formaldehyde fixation for 10 minutes at room temperature. Cells were then washed three times with PBS for five minutes every time. PBS with 0.1% TritonX-100 (PBS_Triton) was used for penetration. BSA of 2% in PBST was used for blocking before staining with antibodies. The first antibodies were diluted by PBST with 1% BSA. Samples were incubated with the first antibodies at 4°C overnight. Then cells were washed three times with 10 minutes for each before being incubated with the secondary antibodies for one hour at room temperature. After antibody incubation, cells were washed with PBS_Triton for five minutes and stained with DAPI (Fisher) for 10 minutes at room temperature. Cells were washed with PBS_Triton for five minutes three times and stored in PBS for imaging.

Photos were taken with Nikon A1 confocal microscopy at CBI. The microscope was controlled by the build-in software, Nikon NIS Elementary. All photos, except the photo for GSK3^AA^ subcellular localization, were taken with plan fluor 40x DIC M/N2 oil objective with 0.75 numerical aperture and 0.72 mm working distance. The scan field were chosen randomly all over the glass-bottom area. For identifying the GSK3^AA^ subcellular localization, plan apo λ 100x oil objective with 1.45 NA and 0.13 mm WD was used. The 3D model of GSK3^AA^ overlapped with ERC and DAPI were reconstructed from 25 of z-stack images in 11.6 μm and videos were produced also by NIS Element software. To minimize photobleaching, an object field was firstly chosen by fast scan, then the photos were taken at 2014 X 2014 pixel or 4096 X 4096 pixel resolution, for generation large data or for photo presentation, respectively.

CellProfiller was used for cell segregation and initial imaging analysis as what described in Carpenter et.al.^50^.

#### Image Correction

To keep identical background through all images, background correction was performed before further image processing. For each image fluorescent intensities in space without cells were used as local background. Photos that have obviously uneven illumination and background fluorescence were removed from further processing. Otherwise the mean background fluorescent intensity was obtained through averaging over the whole image, and was deducted uniformly from the image.

#### Image Segmentation

Cell number and position were determined by nuclear recognition with DAPI. The global strategy was used to identify the nuclear shape, and the Otsu algorithm was used for further calculation. Clumped objectives were identified by shape and divided by intensity. Next, using the shrank nuclear shape as seed, cell shape was identified by the Watershed algorithm. For identifying the clusters of GSK3^AA^ formed around a nucleus, the nuclear shape was shrank manually by 3 pixels and used as a new seed to grown the boundary with the watershed method until reaching background intensity level. All parameters were optimized through an iterative process of automatic segmentation and manual inspection.

#### Image quantification

Averaged fluorescence density and integrated fluorescence intensity were calculated automatically with CellProfiler. The amount of the GSK3^AA^ form was quantified as the sum of intensities of pixels belonging to the cluster formed around a nucleus. Concentrations of all other proteins were quantified by the average pixel intensity within the nucleus or cytosol region of a cell. Next, the quantified results were examined manually, and those cells with either cell area, nuclear area, or fluorescent intensity beyond five folds of the 95% confidence range of samples from a given treatment were discarded, which account for less than 1% of the cells analyzed. Immunofluorescence data were further processed and plots were generated using customized R codes and Matlab codes. The R package^2^, was also used in data analysis.

## Quantitative PCR

Cells were seeded in 12-well plastic bottom cell culture plates and treated as described above. Three parallel experiments were performed in every treatment. Total RNA was isolated with the TRIZOL RNA isolation kit (Fisher), and mRNA was reversely transcribed with the RNAscript II kit (ABI). The stem-loop method was used for microRNA reverse transcription. The qPCR system was prepared with the SYBR green qPCR kit with designed primers (Supplementary table 1) and performed on StepOnePlus real-time PCR (ABI).

## Immunoprecipitation and silver staining

Immunoprecipitation was performed with SureBeads magnetic beads (Bio-Rad) following a protocol modified from the one provided by the manufacture. We washed beads with PBS with 0.1% Tween 20 (PBS_Tween) for three times, then harvested cells by RIP A (Thermo) with proteinase and phosphatase inhibitor (Roche). Samples were pre-cleaned with 100 μl of suspended Protein G per 450 μl of lysis mixture. Antibodies targeting GSK3 (Cell Signaling), GSK3AA (Santa Cruz), and GSK3D (Santa Cruz) were added into every 100 μl of bead mixture respectively. The mixture was rotated at 4 °C for 3 hours. Beads that were conjugated with antibodies were washed with PBS_Tween. An amount of 100 μl of pre-cleaned lysis buffer was added into conjugated beads and rotated at 4 °C overnight. Targeted proteins were eluded from beads by incubating with 40 μl 1x Laemmli buffer with SDS at 70 °C for 10 minutes. For the samples an amount of 5 μl was used for western blot assay, and an amount of 30 μl was loaded for SDS-PAGE (Bio-Rad) and followed by silver staining (Fisher).

## Network reconstruction and coarse-graining

The full network from TGF-β1 to SNAIL1 (Fig. S2A) was generated with IPA (Qiagen®). Specifically, all downstream regulators of TGF-β1 and upstream regulators of SNAIL1 in human, mice and rat were searched and added to the network. Then, direct or indirect relationships between every pair of regulators were searched and added to the network. After obtaining the whole network, regulators that have been reported to be activated later than SNAIL 1 were removed. Examination of the network reveals that the network can be further organized into three groups: the TGF-β-SMAD-SNAIL canonical pathway, the TGF-β-GSK3-β-catenin pathway that has the most number of links, and others. We further noticed that GLI1 is a central connector of TGF-β, SMAD, GSK3 and SNAIL1. We performed western blot studies on β-CATENIN and found that neither its concentration nor its location changes significantly before day 3, therefore we removed β-CATENIN from the network. In addition, previous studies report that the SMAD-GLI axis plays important role in TGF-β induced EMT ^31^. Therefore we further grouped the network as the SMAD module, the GLI module, and the GSK3 module, as well as the remaining ones that we referred as “Others”, and reached the network shown in Fig. 2A. Those molecular species not explicitly specified in Fig. 2A either have their effects implicitly included in the links, for example the link from TGF-β to GSK3, or are included in the links of “Others”. This treatment is justified since our various inhibition experiments indeed showed that the three factors we identified affect SNAIL1 expression the most. These “other” species may contribute to snail1 activation at a time later than what considered in this work. Therefore we emphasize the network in Fig. 2A is valid only within the time window we examined, i.e., within three days after TGF-β1 treatment for MCF10A cells.

## Acknowledgements

This work was supported by the National Science Foundation [DMS-1462049 to JX], and the Pennsylvania Department of Health (SAP 4100062224). We would like to acknowledge the NIH supported microscopy resources in the Center for Biologic Imaging at University of Pittsburgh, specifically the confocal microscope supported by grant number 1S10OD019973-01.

---

**Figure S1.**
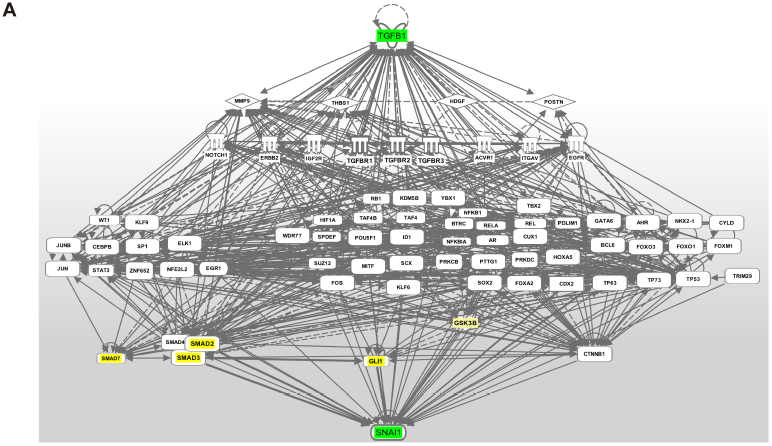
Network of TGF-β activating SNAIL1 reconstructed with IPA.

**Figure S2.**
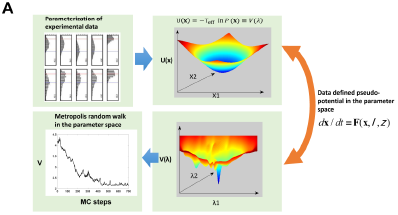
Schematic of the parameter space search approach.

**Figure S3.**
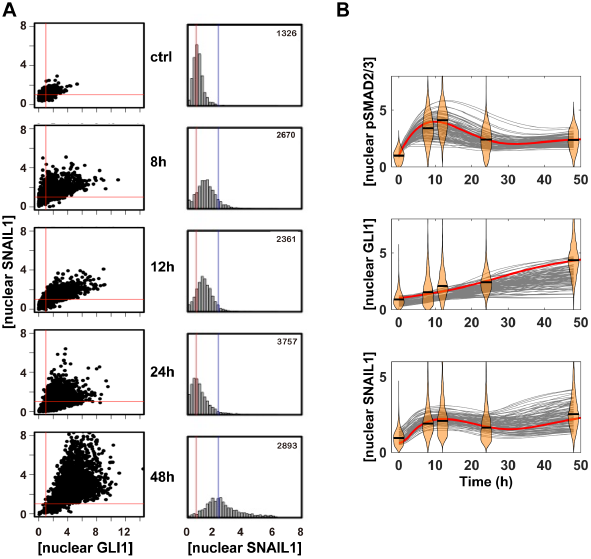
Supplemental results showing GLI1 contributes to the second wave of SNAIL1. (A) Scatted plot of measured nuclear GLI1 and SNAIL1 concentrations and the corresponding histogram representation for [nuclear SNAIL1]. The same sets of data of Fig. 2C are used. (B) The model of Fig. 2A with GLI1 reproduces the observed pSMAD2/3-SNAIL1 dynamics. To fit the SNAIL 1 dynamics the exact temporal profile of GLI1 is not important except the requirement of its activation after 24 h.

**Figure S4.**
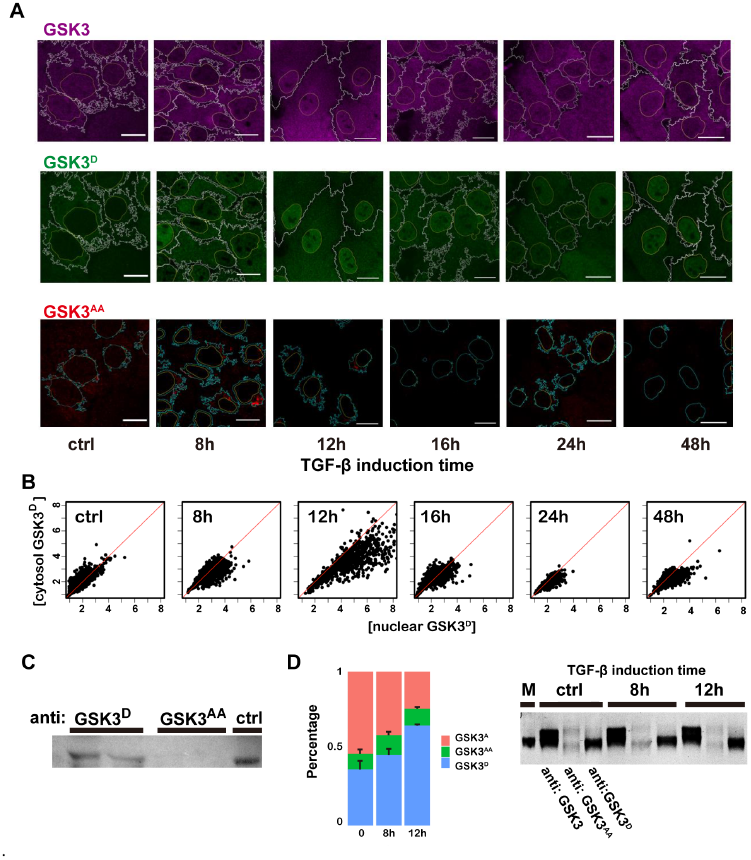
Supplemental results showing temporal switch between two phosphorylation forms of GSK3. (A) IF images showing the temporal switch between two phosphorylation forms of GSK3. (B) Scattered plots showing correlation between nuclear and cytosol concentrations of GSK3^D^. (C) Immunoprecipitation studies showing two phosphorylation forms do not coexist. Data of two replicas was shown. (D) Silver staining measurement of the relative amount of different GSK3 forms. The right figure shows a representative of three independent replicates. M refers to the marker indicating protein mass.

**Figure S5.**
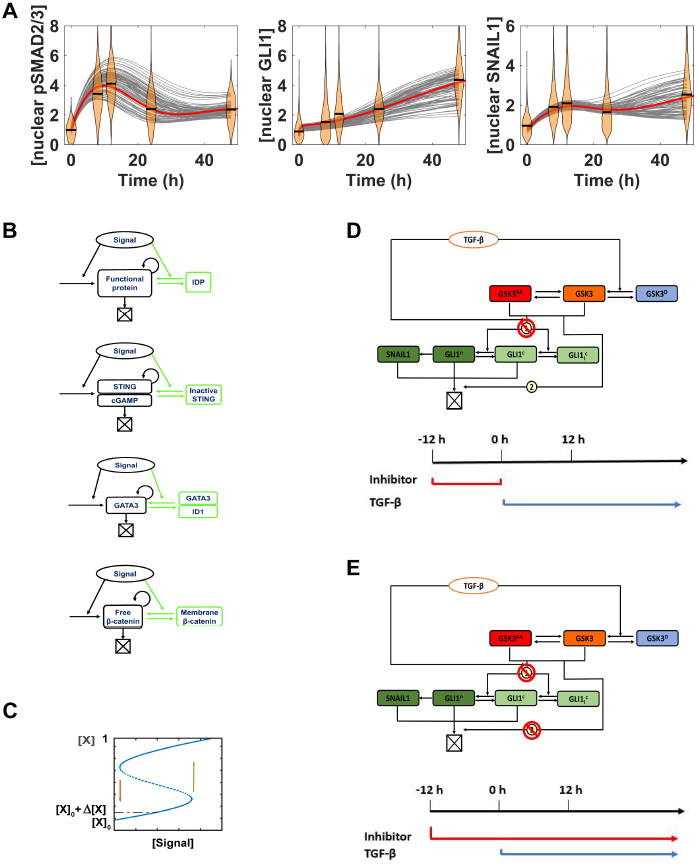
Supplemental results of the full model. (A) The model of Fig. 5A reproduces the observed GLI1 as well as pSMAD2/3-SNAIL 1 dynamics. (B) Examples of regulatory factors having positive feedback loop and reservoir of molecules in inactive form that can be activated by another stimulus. IDPs refer to intrinsically disordered proteins, and some of them are transcription factors, which change into folded form and have higher DNA binding affinity upon binding of cofactors or posttranslational modification. ID1 is a member of the family of inhibitors of DNA binding proteins. (C) Bifurcation diagram showing that the initial concentration boost is small compared to the concentration jump associated with external signal induced switch of cell states. (D-E) Schematics of the early and full GSK3 inhibition experiments.

**Figure S6.**
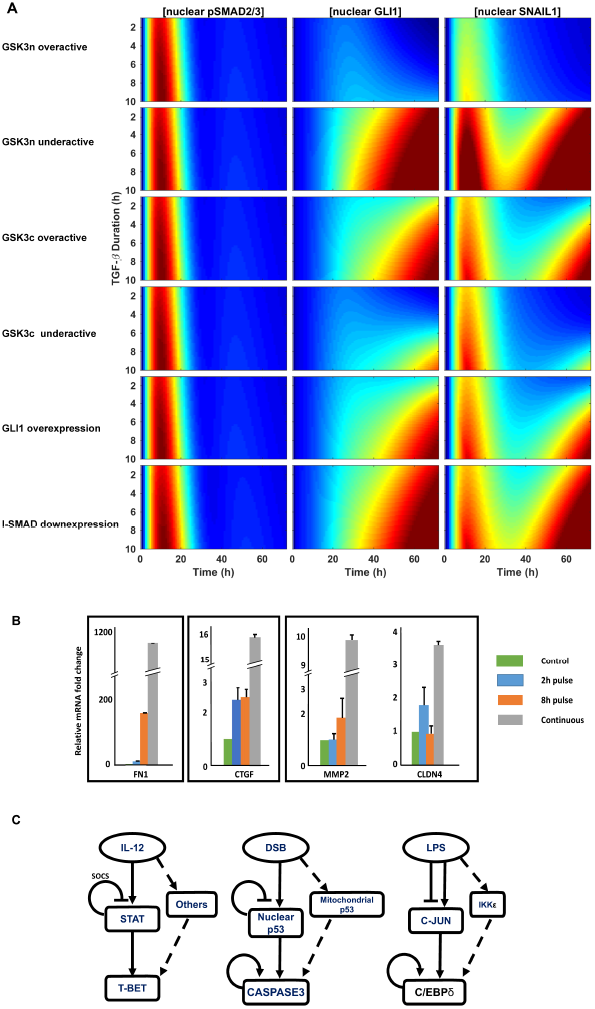
Supplemental results that cells can detect signal duration. (A) Supplemental model results of pulsed TGF-β1 treatments with various mutations. (B) The mRNA levels of selected TGF-β activated genes at day 3 after different durations of TGF-β1 treatments. (C) Examples of other signaling transduction pathways that share similar motif structure as that of TGF-β signaling, including IL-12, DNA double strand breaking, and LPS.

**Supplementary Movie S1: Subcellular localization of GSK3^AA^ (red).** Movies were composed from z-stack imaging.

**Supplementary Movie S2: Subcellular localization of GSK3^AA^ (red) overlaid with ERC (green) and DAPI (blue, nuclear area).** Movies were composed from z-stack imaging.

## Supplementary Materials

### Mathematical modeling

#### Canonical TGF-β/pSMAD2/3/SNAILl pathway (Fig. 1A)

We used this ordinary differential equation (ODE) model in Fig. 1 and Fig. S1.

##### TGF-β/SMAD2/3 module

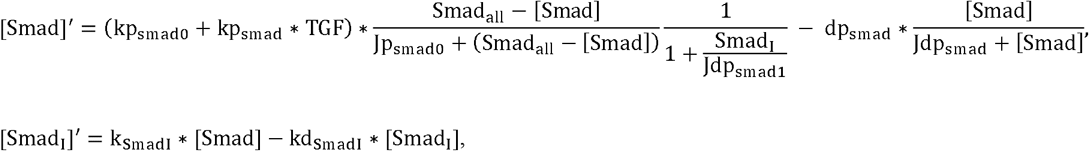

where [Smad] and [Smadj] are the concentrations of pSMAD2/3 and inhibitory SMAD, respectively.

##### SNAIL-miR-34 module

It is expanded from our previous model^33^ by considering transcription activation of SNAIL1 by pSMAD2/3 and TGF-β, and degradation of SNAIL1.

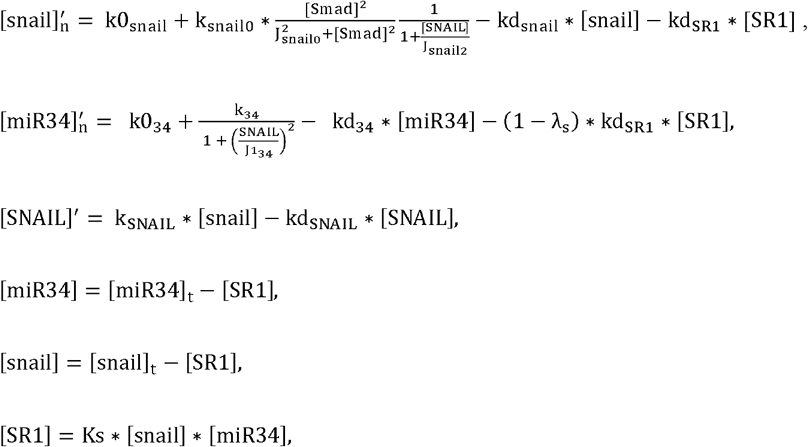

where [snail], [miR34], [SNAIL], [snail], [SRI] are the concentrations of total *SNAIL1* mRNA, miR-34, SNAIL 1 protein, free *SNAIL1* mRNA and *miR-34-SNAIL1* mRNA complex, respectively.

#### Canonical TGF-β/SMAD/SNAILl pathway with GLI1 (Fig. S2)

Taking into account the GLI1 self-activation and GLI1 mediated expression of *SNAIL1* mRNA, we added another ODE for GLI1 and revised the ODE of *SNAIL1* mRNA.

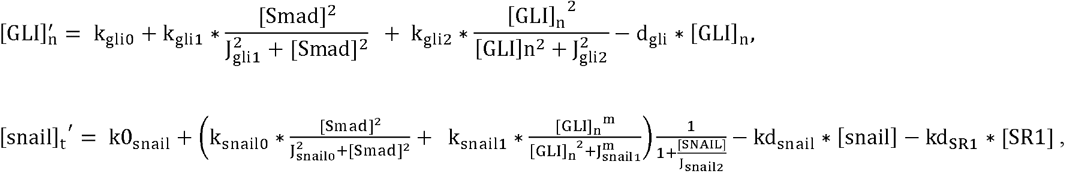

where [GLI]_n_ is the concentration of nuclear GLI1. We used this ODE model to generate results in Fig. S2.

#### Model for the GSK3/GLI module (Fig. 4A)

Since the process involves many steps and a detailed model would require many parameters to determine, instead we used two phenomenological time-dependent functions to qualitatively mimic the dynamics of the enzyme activities of cytosol GSK3 and nuclear GSK3 we experimentally measured,

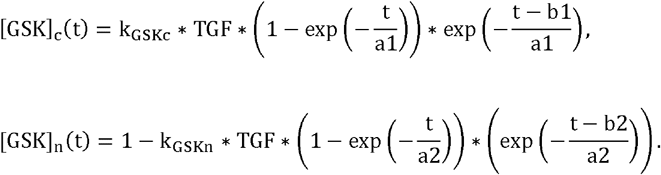

Furthermore, the basal pool of cytosol GLI1 is considered, which is by sequestered in the cytosol by Sufu but could translocate to the nuclear after Sufu is inactivated by the cytosol enzyme GSK3 activity. We used a revised ODE of nuclear GLI1 concentration derived with the quasi-equilibrium approximation (see below)

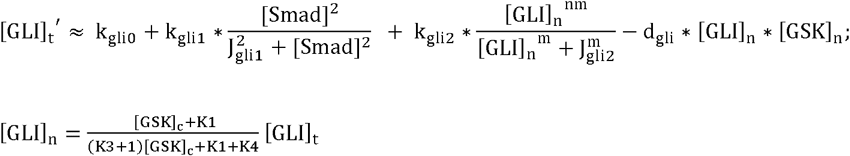

##### Derivation ofGLI ODE

We assumed the quasi-equilibrium approximation for the GLI nuclear and cytosol shuttling, the GSK3 regulated binding/unbinding between Sufu and GLI in the cytosol, and obtained the following equations,

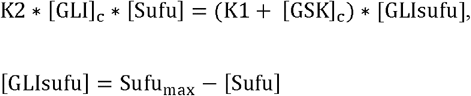

Thus we have

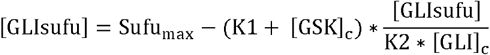

That is,

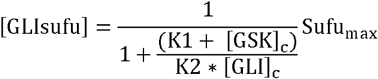

Also we have

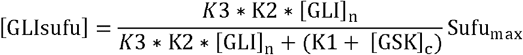

thus

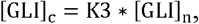

The total level of GLI1 is the sum of the three forms, **GLISufu, GLI_C_** and **GLI_n_,**

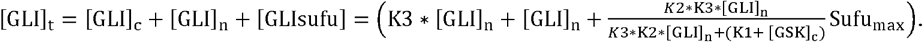

Thus, we obtained the relation among ***[GLI]_n_, [GLI]_t_***and **[GSK]_C_**

[GLI]_n_=*f*(*GSK*_c_,[GLI]_t_),

The total concentration of GLI1 is given by,

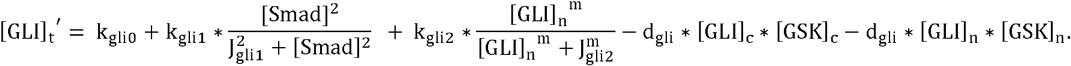

Given that our data shows that **[GLI]_C_** is low throughout the process, we neglected the degradation term of **[GLI]_C_,**

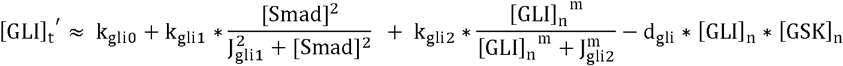

##### TGF-βpulse

Since TGF-β1 can enter to cells through endocytosis, washing the extracellular TGF-β1 does not stop the signaling immediately. Therefore, we modeled the effective TGF-β1 concentration by the following equation,

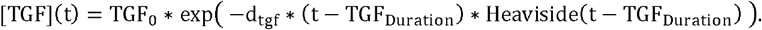

##### Full model

By considering all the modules, the full model is as following,

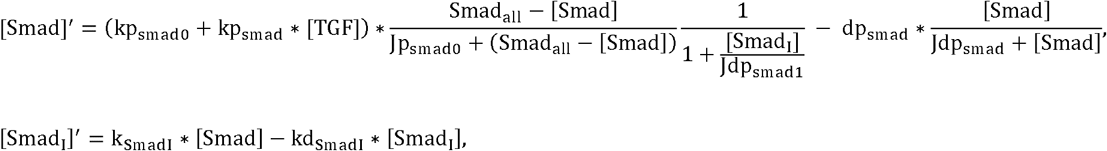

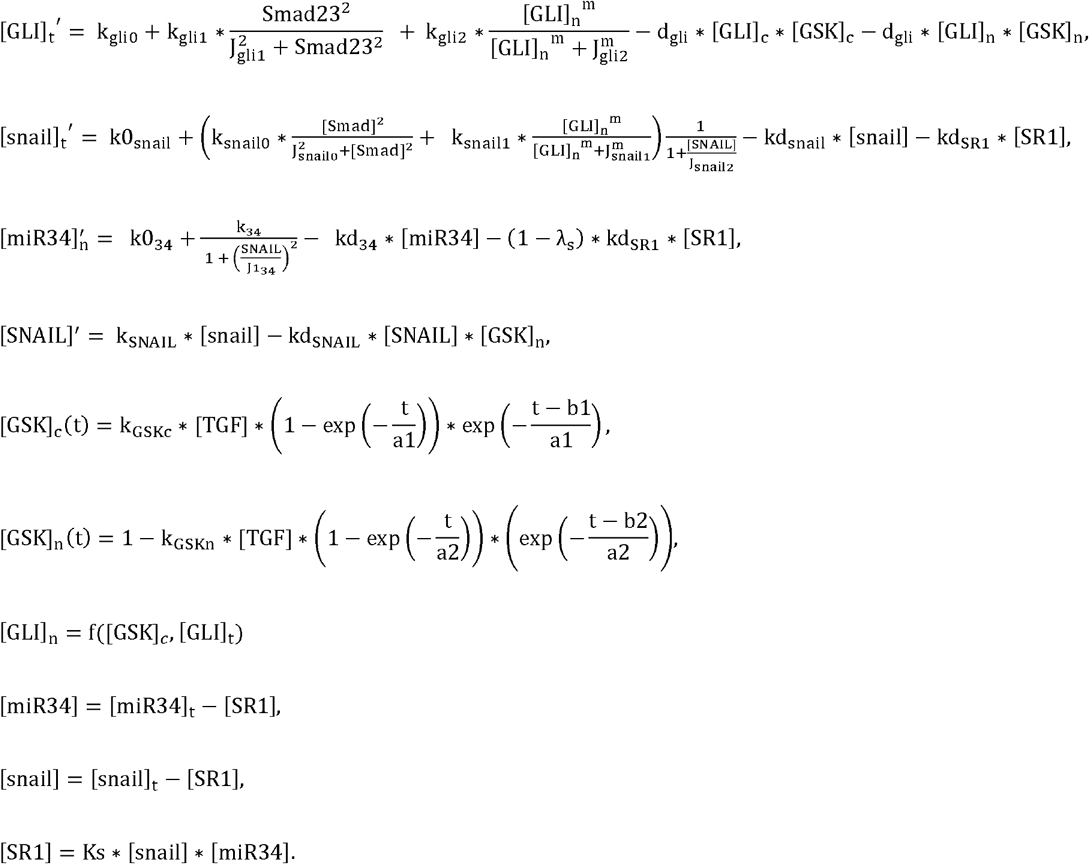

We used this ODE model to generate results in Fig. 5 and Fig. S5 A.

##### Parameter space searching

###### Step 1: Calculate single cell distributions of experimental observables

We calculated histograms of the distributions from the single cell experimental data. Suppose that we have *N* observables measured in *M* time points, we have an *N* × *M* dimensional distribution of the data. Since we used fixed cells and we had no information on the temporal correlation, we treated the distributions from different time points as independent, *i.e.,* 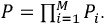.

###### Step 2: Define pseudo-potentials from the parameterized distribution

We defined a pseudo-scalar-potential function *U*(**x_1_, **x**_2_,.., **x**_M_) = −*T*_*eff*_**(***inP****−*lnP*_*max*_).* The constant *T_eff_* is an effective temperature, which we chose *T_eff_* = 1. The constant term *lnPmax* sets the potential to be zero at the peak position of the distribution, and does not affect the parameter space search results. This pseudo-potential is just an auxiliary scalar function for the following application of the Metropolis algorithm. If a mathematical model can faithfully describe the system dynamics, with given initial conditionals and non-adjustable parameter set of ζ, we should be able to find distributions of the parameter set *λ* (to take into account cell-to-cell heterogeneity), and generate the corresponding distributions of *(x_ll_x_2_,*. .,x_M_) to reproduce *U.* That is, for a specific set of *λ,* **x**_i_ = **X_i_**(**x**_0_; **λ**, **ζ**),i = 1, … *M*, and *U*(**x_1_, x_2_,.., x_M_**) = *V(λ).* Unlike *U,* the function form of *V* can be very complex, but fortunately we do not need to know its explicit function form to perform the following Metropolis sampling.

###### Step 3: Obtain model parameter distributions that reproduce the distributions of experimental observables

Now it is clear why we define the pseudo-potential. We performed Monte Carlo random walks along the pseudo-potential Vin the *λ* space using the Metropolis algorithm, just as how the algorithm is typically applied along real physical potentials. At each step with a set of *λ,* we generated a trial move *λ’ = λ* + *δλ.* We propagated the ODEs to obtain *V(λ)* and *V(λ’),* then use the Metropolis criteria to decide whether to accept the new move. If *V*(λ’) ≤ *V*(*λ*) accept this step and update the parameter set **λ** = λ’. If *V*(λ’) > *V*(λ) , accept this step with a probability exp(−(*V*(λ’) − *V*(λ))/*T*), with *T* = 1.

In our model, there is no feedback between the SMAD2/3 module and the SNAIL1/miR-34 module, thus we used a two-step to searching the parameter space for the TGF-β/SMAD2/3 module,

1. Search the parameter space (nine parameters) in the SMAD2/3 module;
2. Search the parameter space (six parameters) for the SNAIL1/miR-34 module based on the 50 samples of good-fit parameter set of the SMAD2/3 module from step 1.

In step 2 some of the parameters in the SNAIL1/miR-34 module were fixed and used as a well-trained parameter set from our previous work ^33^. Instead only six new parameters that connect the module SMAD2/3 and module SNAIL1/miR-34 were considered in the parameter space searching.

When the GLI1 module was included, we again used the fact that there is no feedback between the SMAD2/3 module and the GLI1 module, and used a three-step searching procedure to reduce the computational efforts,

1. Search the parameter space (nine parameters) for the SMAD2/3 module;
2. Search the parameter space (seven parameters) for the GLI1 module;
3. Search the parameter space (six parameters) for the SNAIL1/miR-34 module based on the 50 samples of good-fit parameter set of the SMAD2/3 module the GLI1 module from step 1-2.

#### Parameter change in various over-expression/down-expression or over-active/down-active conditions (Fig. S5)

To produce the results in Fig. S5A, a 1.2-fold change of *k_glio_* is used in the case of GLI1 over-expression, a 0.8-fold change of *k_gskn_* in the case of I-SMAD down-regulation. There is 0.8-fold change of *k_gskn_* in the case of over-active cytosol GSK3, 1.2-fold change of *k_gskn_in* the case of under-active cytosol GSK3. Similarly, there is 1.2-fold change of *k_gskc_* in the case of over-active nuclear GSK3, and 0.5-fold change of *k_gskc_* in the case of under-active nuclear GSK3.

**Supplementary Table 1.**
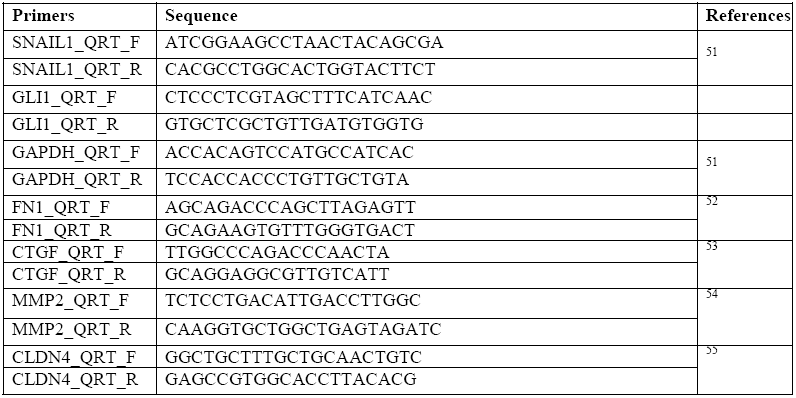
Primer list.

**Supplementary Table 2.**
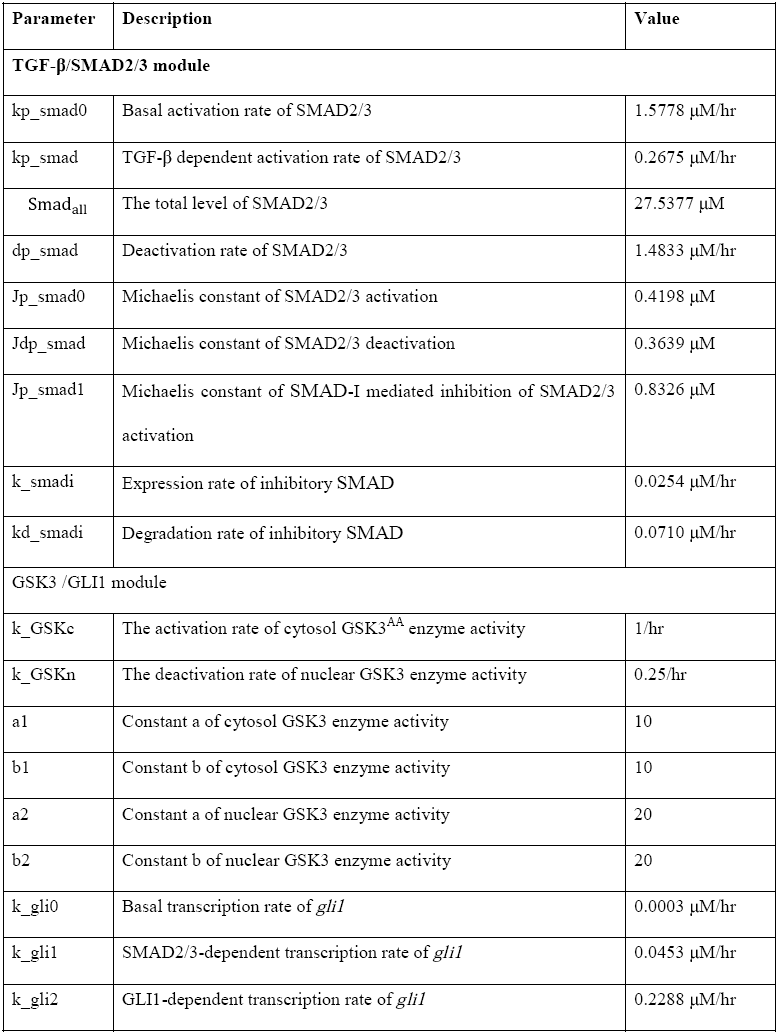
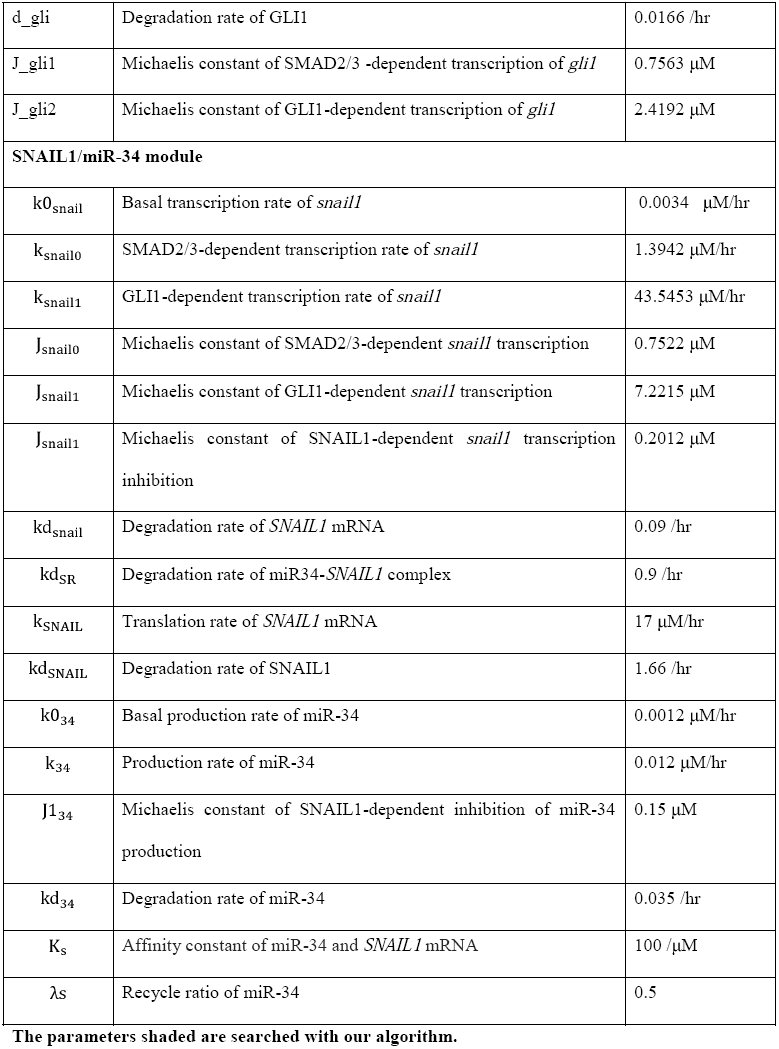
The Parameters values of the best fit of the full model.

The parameters shaded are searched with our algorithm.

